# Intake of red and processed meat, use of non-steroid anti inflammatory drugs, genetic variants and risk of colorectal cancer; a prospective study of the Danish “Diet, Cancer and Health” cohort

**DOI:** 10.1101/496968

**Authors:** Vibeke Andersen, Ulrich Halekoh, Anne Tjønneland, Ulla Vogel, Tine Iskov Kopp

## Abstract

Red and processed meat have been associated with increased risk of colorectal cancer (CRC), whereas long-term use of non-steroid anti-inflammatory drugs (NSAIDs) may reduce the risk. The aim was to investigate potential interactions between meat intake, NSAID use, and gene variants in fatty acid metabolism and NSAID pathways in relation to the risk of CRC. A nested case-cohort study of 1038 CRC cases and 1857 randomly selected participants from the Danish prospective “Diet, Cancer and Health” study encompassing 57,053 persons was performed using the Cox proportional hazard models. Gene variants in *SLC25A20, PRKAB1, LPCAT1, PLA2G4A, ALOX5, PTGER3, TP53, CCAT2, TCF7L2, BCL2* were investigated. *CCAT2* rs6983267 was associated with risk of CRC per se (p<0.01). Statistically significant interactions were found between intake of red and processed meat and *CCAT2* rs6983267, *TP53* rs1042522, *LPCAT1* rs7737692, *SLC25A20* rs7623023 (p_interaction_=0.04, 0.04, 0.02, 0.03, respectively), and use of NSAID and alcohol intake and *TP53* rs1042522 (p_interaction_=0.04, 0.04, respectively) in relation to risk of CRC. No other consistent associations or interactions were found. This study replicated an association of *CCAT2* rs6983267 with CRC and an interaction between *TP53* rs1042522 and NSAID in relation to CRC. Interactions between genetic variants in fatty acid metabolism and NSAID pathway and intake of red and processed meat were found. Our results suggest that meat intake and NSAID use affect the same carcinogenic mechanisms. All new findings should be sought replicated in independent prospective studies. Future studies on the cancer-protective effects of aspirin/NSAID should include gene and meat assessments.

**Author Summary:** Intake of red and processed meat has been associated with risk of cancer and in particular colorectal cancer. However, the underlying biological mechanisms are only incompletely understood. Gene-environment interaction analysis may be used for identifying underlying mechanisms for e.g. meat carcinogenesis. In this work, we have analyzed the interaction between the intake of red and processed meat, use of non-steroid anti-inflammatory drugs (including the anti-carcinogenic drug aspirin) and genetic variants. Our results suggest that meat intake and non-steroid anti-inflammatory drug use affect the same carcinogenic mechanisms. These results need to be replicated in other cohort studies with lifestyle information. If replicated, these results may have future implications for developing new strategies for preventing colorectal cancer and other cancers that share similar pathways.

## Introduction

Colorectal cancer (CRC) is the third most common malignant tumor and the fourth leading cause of cancer death worldwide with a lifetime risk in Western European and North American populations of around 5% [1]. Multiple risk factors, both genetic and environmental, are involved in the etiology and prognosis of CRC [2]. Identification and characterization of the risk factors, their potential interactions, and the underlying biological mechanisms are requested as a basis for improving preventative strategies that may include identifying individuals who would most benefit from preventive strategies.

Epidemiological studies suggest that high intake of red and particularly processed meat may increase the CRC risk [3], whereas long-term use of non-steroid anti-inflammatory drugs (NSAIDs) including aspirin (acetylic acid) may reduce the risk of CRC [4, 5]. Investigations on the potential carcinogenic mechanisms of red and processed meat have suggested that meat may confer carcinogenesis by being a source of cooking mutations (heterocyclic amine, *N*-nitroso compounds) formed during preparation [6], organic sulfur-containing proteins leading to a high content of H_2_S in the intestinal lumen, a highly potent regulator of intestinal cell function including inflammation and cell death signaling [7] and/or microbial factors arising during storage [8]. Similarly, the underlying cancer protective mechanisms of NSAID have been investigated and both COX-2 dependent and COX-2 independent mechanisms have been suggested [9, 10].

Still, however, the mechanisms are incompletely understood. First of all, epidemiological studies are not suitable to evaluate CRC causality because of colinearity between the studied factors (intake of red and processed meat and NSAID) and other potential CRC risk factors (such as e.g. Western-style diet and high body mass index) that limit the ability to analytically isolate the independent effects of the studied factors [11]. Next, although animal studies may suggest important underlying biological mechanisms [12], results from animal studies may not apply to humans due to differences in the biology such as the metabolism of meat between animals and humans and because doses used in animals may not be transferable to human conditions [6]. Gene-environment (GxE) interaction analyses may overcome the methodological issues mentioned above. Indeed, the identification of an interaction between a genetic variant (functional or in linkage with a functional variant) in a gene that is chosen based on its biological function and an environmental factor suggests that both factors are involved in the same process. Using GxE interaction analysis, we have investigated potential mechanisms by which red and processed meat and NSAID may affect CRC carcinogenesis [13-21] (reviewed in [22-24]). Red and processed meat is a rich source of n-6 polyunsaturated fat that is converted into arachidonic acid after ingestion and further metabolized into several bioactive lipids that play critical roles in a variety of biologic processes involved in chronic inflammation and colorectal cancer. Conversely, NSAIDs including aspirin may reduce inflammation and CRC risk via similar and other pathways in relation to CRC [22, 23, 25].

Thus, the aim of the present study was to investigate the association of polymorphisms in genes involved in fatty acid metabolism and NSAID pathway with CRC, and, furthermore, interactions between these polymorphisms and NSAID use and dietary factors focusing on the intake of red and processed meat in relation to CRC. The study cohort was the Danish “Diet, Cancer and Health” with prospectively collected lifestyle information encompassing 57053 participants whereof 1038 cases that developed CRC were compared to a sub-cohort of 1857 members using a nested case-cohort design. In addition to replicating earlier findings, this study found interactions between genetic variants in fatty acid metabolism and NSAID pathway and intake of red and processed meat suggesting that meat intake and NSAID use affect the same carcinogenic mechanisms.

## Results

**Table 1** shows the baseline characteristics of 1038 CRC cases and 1857 sub-cohort members including CRC risk factors. Among the controls, the genotype distributions of the studied polymorphisms were in Hardy–Weinberg equilibrium (results not shown). In order to maximize the statistical power for the interactions analyses, the genotypes were combined assuming either a dominant model (*SLC25A20* rs7623023, *TP53* rs1042522, *CCAT2* rs6983267, *BCL2* rs2279115) or a recessive model (*PRKAB1* rs4213, *LPCAT1* rs7737692, *PLA2G4A* rs4402086, *ALOX5* rs3780894, *PTGER3* rs6685546, *TCF7L2* rs7903146) based on the observed risk estimates.

**Table 1.**
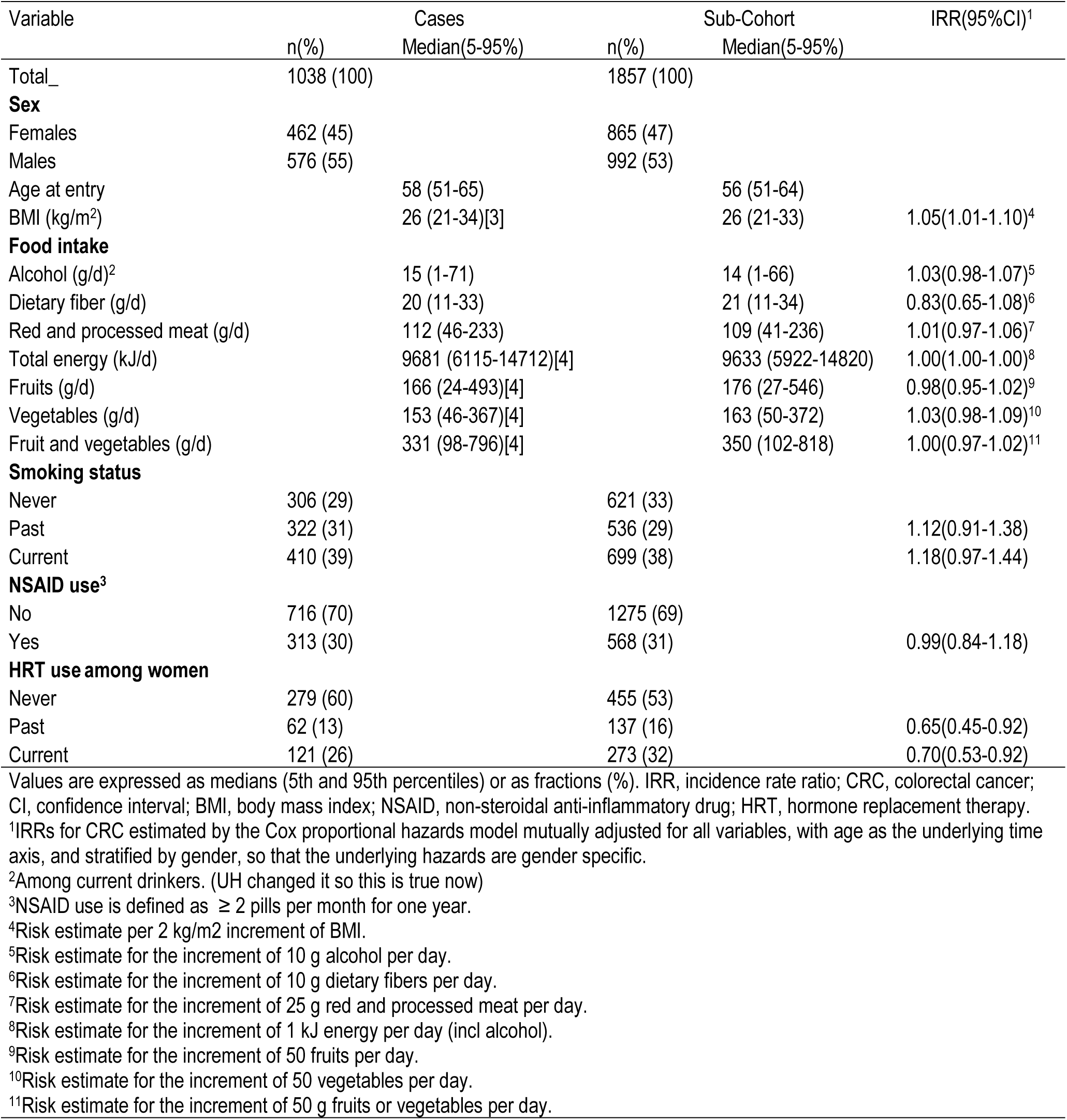
Participant description.

| Variable | Cases | Sub-Cohort | IRR(95%CI)<sup>1</sup> |
| --- | --- | --- | --- |
|  | n(%)<br>Median(5-95%) | n(%)<br>Median(5-95%) |  |
| Total_ | 1038 (100) | 1857 (100) |  |
| **Sex** |  |  |  |
| Females | 462 (45) | 865 (47) |  |
| Males | 576 (55) | 992 (53) |  |
| Age at entry | 58 (51-65) | 56 (51-64) |  |
| BMI (kg/m<sup>2</sup>) | 26 (21-34)[3] | 26 (21-33) | 1.05(1.01-1.10)<sup>4</sup> |
| **Food intake** |  |  |  |
| Alcohol (g/d)<sup>2</sup> | 15 (1-71) | 14 (1-66) | 1.03(0.98-1.07)<sup>5</sup> |
| Dietary fiber (g/d) | 20 (11-33) | 21 (11-34) | 0.83(0.65-1.08)<sup>6</sup> |
| Red and processed meat (g/d) | 112 (46-233) | 109 (41-236) | 1.01(0.97-1.06)<sup>7</sup> |
| Total energy (kJ/d) | 9681 (6115-14712)[4] | 9633 (5922-14820) | 1.00(1.00-1.00)<sup>8</sup> |
| Fruits (g/d) | 166 (24-493)[4] | 176 (27-546) | 0.98(0.95-1.02)<sup>9</sup> |
| Vegetables (g/d) | 153 (46-367)[4] | 163 (50-372) | 1.03(0.98-1.09)<sup>10</sup> |
| Fruit and vegetables (g/d) | 331 (98-796)[4] | 350 (102-818) | 1.00(0.97-1.02)<sup>11</sup> |
| **Smoking status** |  |  |  |
| Never | 306 (29) | 621 (33) |  |
| Past | 322 (31) | 536 (29) | 1.12(0.91-1.38) |
| Current | 410 (39) | 699 (38) | 1.18(0.97-1.44) |
| **NSAID use<sup>3</sup>** |  |  |  |
| No | 716 (70) | 1275 (69) |  |
| Yes | 313 (30) | 568 (31) | 0.99(0.84-1.18) |
| **HRT use among women** |  |  |  |
| Never | 279 (60) | 455 (53) |  |
| Past | 62 (13) | 137 (16) | 0.65(0.45-0.92) |
| Current | 121 (26) | 273 (32) | 0.70(0.53-0.92) |

Values are expressed as medians (5th and 95th percentiles) or as fractions (%). IRR, incidence rate ratio; CRC, colorectal cancer; CI, confidence interval; BMI, body mass index; NSAID, non-steroidal anti-inflammatory drug; HRT, hormone replacement therapy.
<sup>1</sup>IRRs for CRC estimated by the Cox proportional hazards model mutually adjusted for all variables, with age as the underlying time axis, and stratified by gender, so that the underlying hazards are gender specific.
<sup>2</sup>Among current drinkers. (UH changed it so this is true now)
<sup>3</sup>NSAID use is defined as $\geq 2$ pills per month for one year.
<sup>4</sup>Risk estimate per 2 kg/m<sup>2</sup> increment of BMI.
<sup>5</sup>Risk estimate for the increment of 10 g alcohol per day.
<sup>6</sup>Risk estimate for the increment of 10 g dietary fibers per day.
<sup>7</sup>Risk estimate for the increment of 25 g red and processed meat per day.
<sup>8</sup>Risk estimate for the increment of 1 kJ energy per day (incl alcohol).
<sup>9</sup>Risk estimate for the increment of 50 fruits per day.
<sup>10</sup>Risk estimate for the increment of 50 vegetables per day.
<sup>11</sup>Risk estimate for the increment of 50 g fruits or vegetables per day.

### Associations between polymorphisms and CRC

**Table 2** shows the crude associations between the SNPs and CRC. There was an association between *CCAT2* rs6983267 and CRC (p<0.01). Carriers of the *CCAT2* rs6983267 variant T-allele have about 30% lower risk of CRC compared to GG homozygotes. No other statistically significant associations were found.

**Table 2.**
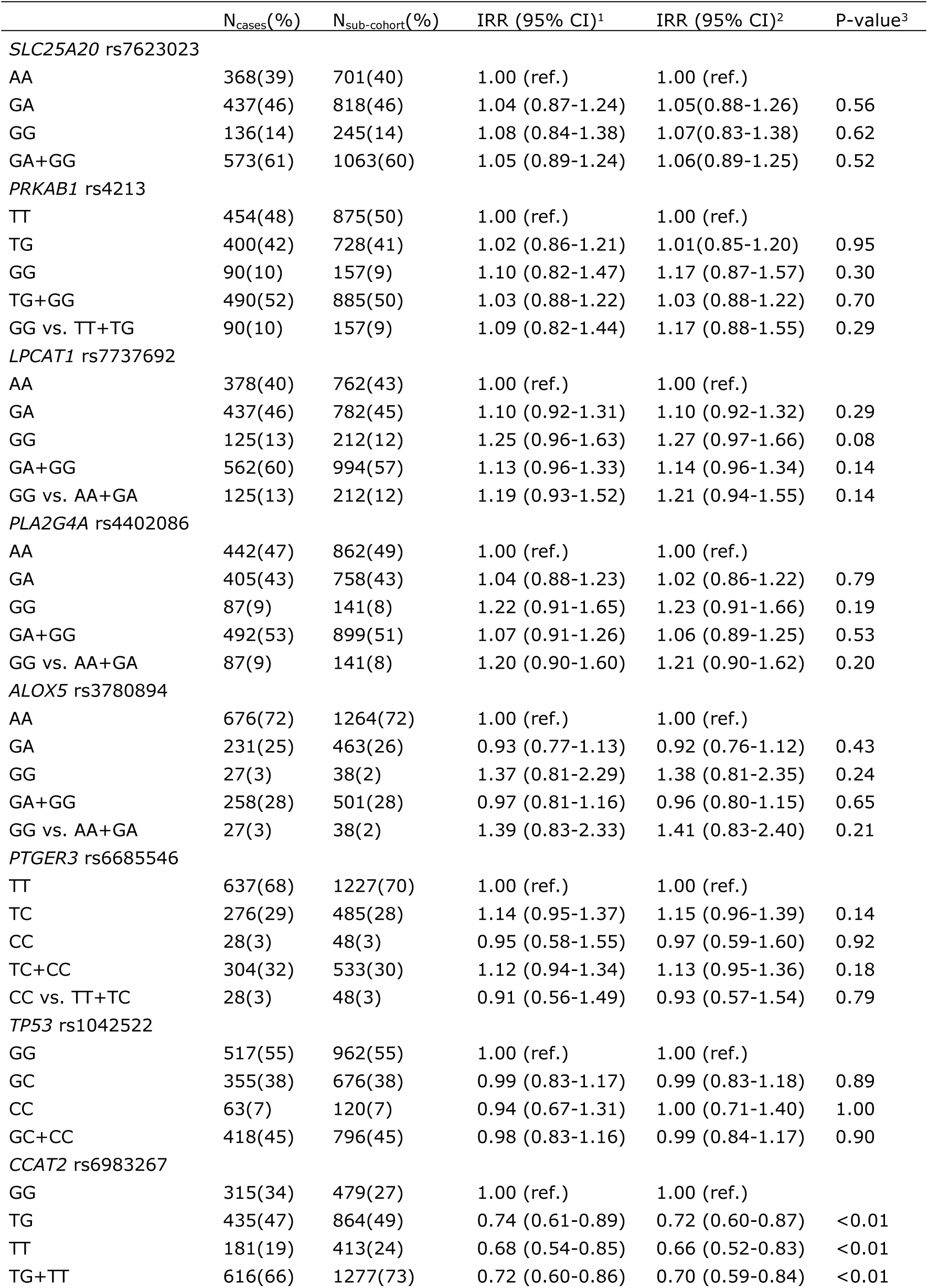
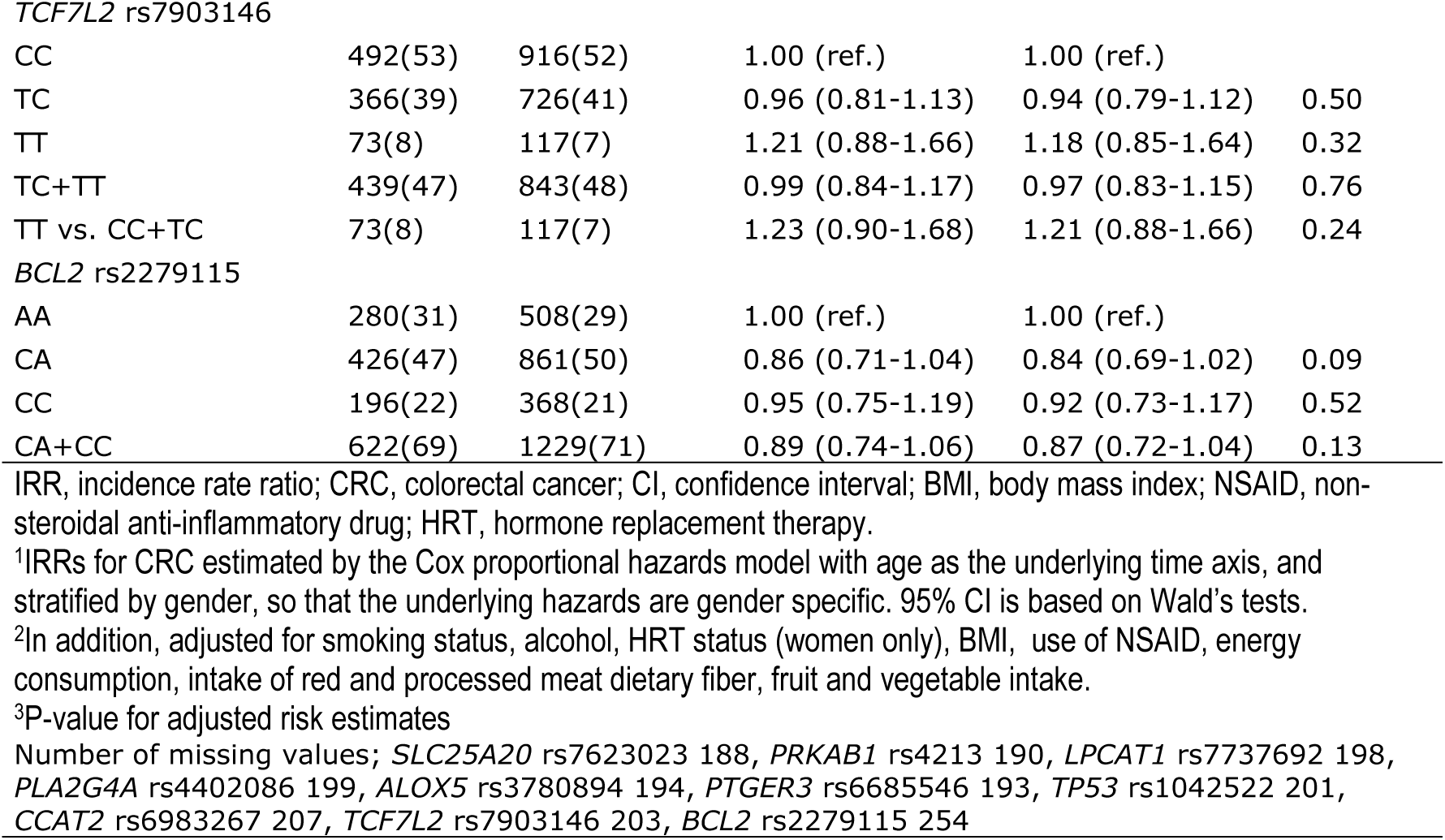
Incidence rate ratios (IRR) for associations with colorectal cancer (CRC)

|  | N<sub>cases</sub>(%) | N<sub>sub-cohort</sub>(%) | IRR (95% CI)<sup>1</sup> | IRR (95% CI)<sup>2</sup> | P-value<sup>3</sup> |
| --- | --- | --- | --- | --- | --- |
| *SLC25A20* rs7623023 |  |  |  |  |  |
| AA | 368(39) | 701(40) | 1.00 (ref.) | 1.00 (ref.) |  |
| GA | 437(46) | 818(46) | 1.04 (0.87-1.24) | 1.05(0.88-1.26) | 0.56 |
| GG | 136(14) | 245(14) | 1.08 (0.84-1.38) | 1.07(0.83-1.38) | 0.62 |
| GA+GG | 573(61) | 1063(60) | 1.05 (0.89-1.24) | 1.06(0.89-1.25) | 0.52 |
| *PRKAB1* rs4213 |  |  |  |  |  |
| TT | 454(48) | 875(50) | 1.00 (ref.) | 1.00 (ref.) |  |
| TG | 400(42) | 728(41) | 1.02 (0.86-1.21) | 1.01(0.85-1.20) | 0.95 |
| GG | 90(10) | 157(9) | 1.10 (0.82-1.47) | 1.17 (0.87-1.57) | 0.30 |
| TG+GG | 490(52) | 885(50) | 1.03 (0.88-1.22) | 1.03 (0.88-1.22) | 0.70 |
| GG vs. TT+TG | 90(10) | 157(9) | 1.09 (0.82-1.44) | 1.17 (0.88-1.55) | 0.29 |
| *LPCAT1* rs7737692 |  |  |  |  |  |
| AA | 378(40) | 762(43) | 1.00 (ref.) | 1.00 (ref.) |  |
| GA | 437(46) | 782(45) | 1.10 (0.92-1.31) | 1.10 (0.92-1.32) | 0.29 |
| GG | 125(13) | 212(12) | 1.25 (0.96-1.63) | 1.27 (0.97-1.66) | 0.08 |
| GA+GG | 562(60) | 994(57) | 1.13 (0.96-1.33) | 1.14 (0.96-1.34) | 0.14 |
| GG vs. AA+GA | 125(13) | 212(12) | 1.19 (0.93-1.52) | 1.21 (0.94-1.55) | 0.14 |
| *PLA2G4A* rs4402086 |  |  |  |  |  |
| AA | 442(47) | 862(49) | 1.00 (ref.) | 1.00 (ref.) |  |
| GA | 405(43) | 758(43) | 1.04 (0.88-1.23) | 1.02 (0.86-1.22) | 0.79 |
| GG | 87(9) | 141(8) | 1.22 (0.91-1.65) | 1.23 (0.91-1.66) | 0.19 |
| GA+GG | 492(53) | 899(51) | 1.07 (0.91-1.26) | 1.06 (0.89-1.25) | 0.53 |
| GG vs. AA+GA | 87(9) | 141(8) | 1.20 (0.90-1.60) | 1.21 (0.90-1.62) | 0.20 |
| *ALOX5* rs3780894 |  |  |  |  |  |
| AA | 676(72) | 1264(72) | 1.00 (ref.) | 1.00 (ref.) |  |
| GA | 231(25) | 463(26) | 0.93 (0.77-1.13) | 0.92 (0.76-1.12) | 0.43 |
| GG | 27(3) | 38(2) | 1.37 (0.81-2.29) | 1.38 (0.81-2.35) | 0.24 |
| GA+GG | 258(28) | 501(28) | 0.97 (0.81-1.16) | 0.96 (0.80-1.15) | 0.65 |
| GG vs. AA+GA | 27(3) | 38(2) | 1.39 (0.83-2.33) | 1.41 (0.83-2.40) | 0.21 |
| *PTGER3* rs6685546 |  |  |  |  |  |
| TT | 637(68) | 1227(70) | 1.00 (ref.) | 1.00 (ref.) |  |
| TC | 276(29) | 485(28) | 1.14 (0.95-1.37) | 1.15 (0.96-1.39) | 0.14 |
| CC | 28(3) | 48(3) | 0.95 (0.58-1.55) | 0.97 (0.59-1.60) | 0.92 |
| TC+CC | 304(32) | 533(30) | 1.12 (0.94-1.34) | 1.13 (0.95-1.36) | 0.18 |
| CC vs. TT+TC | 28(3) | 48(3) | 0.91 (0.56-1.49) | 0.93 (0.57-1.54) | 0.79 |
| *TP53* rs1042522 |  |  |  |  |  |
| GG | 517(55) | 962(55) | 1.00 (ref.) | 1.00 (ref.) |  |
| GC | 355(38) | 676(38) | 0.99 (0.83-1.17) | 0.99 (0.83-1.18) | 0.89 |
| CC | 63(7) | 120(7) | 0.94 (0.67-1.31) | 1.00 (0.71-1.40) | 1.00 |
| GC+CC | 418(45) | 796(45) | 0.98 (0.83-1.16) | 0.99 (0.84-1.17) | 0.90 |
| *CCAT2* rs6983267 |  |  |  |  |  |
| GG | 315(34) | 479(27) | 1.00 (ref.) | 1.00 (ref.) |  |
| TG | 435(47) | 864(49) | 0.74 (0.61-0.89) | 0.72 (0.60-0.87) | <0.01 |
| TT | 181(19) | 413(24) | 0.68 (0.54-0.85) | 0.66 (0.52-0.83) | <0.01 |
| TG+TT | 616(66) | 1277(73) | 0.72 (0.60-0.86) | 0.70 (0.59-0.84) | <0.01 |
| *TCF7L2* rs7903146 |  |  |  |  |  |
| CC | 492(53) | 916(52) | 1.00 (ref.) | 1.00 (ref.) |  |
| TC | 366(39) | 726(41) | 0.96 (0.81-1.13) | 0.94 (0.79-1.12) | 0.50 |
| TT | 73(8) | 117(7) | 1.21 (0.88-1.66) | 1.18 (0.85-1.64) | 0.32 |
| TC+TT | 439(47) | 843(48) | 0.99 (0.84-1.17) | 0.97 (0.83-1.15) | 0.76 |
| TT vs. CC+TC | 73(8) | 117(7) | 1.23 (0.90-1.68) | 1.21 (0.88-1.66) | 0.24 |
| *BCL2* rs2279115 |  |  |  |  |  |
| AA | 280(31) | 508(29) | 1.00 (ref.) | 1.00 (ref.) |  |
| CA | 426(47) | 861(50) | 0.86 (0.71-1.04) | 0.84 (0.69-1.02) | 0.09 |
| CC | 196(22) | 368(21) | 0.95 (0.75-1.19) | 0.92 (0.73-1.17) | 0.52 |
| CA+CC | 622(69) | 1229(71) | 0.89 (0.74-1.06) | 0.87 (0.72-1.04) | 0.13 |

IRR, incidence rate ratio; CRC, colorectal cancer; CI, confidence interval; BMI, body mass index; NSAID, non-steroidal anti-inflammatory drug; HRT, hormone replacement therapy.
<sup>1</sup>IRRs for CRC estimated by the Cox proportional hazards model with age as the underlying time axis, and stratified by gender, so that the underlying hazards are gender specific. 95% CI is based on Wald's tests.
<sup>2</sup>In addition, adjusted for smoking status, alcohol, HRT status (women only), BMI, use of NSAID, energy consumption, intake of red and processed meat dietary fiber, fruit and vegetable intake.
<sup>3</sup>P-value for adjusted risk estimates
Number of missing values; *SLC25A20* rs7623023 188, *PRKAB1* rs4213 190, *LPCAT1* rs7737692 198, *PLA2G4A* rs4402086 199, *ALOX5* rs3780894 194, *PTGER3* rs6685546 193, *TP53* rs1042522 201, *CCAT2* rs6983267 207, *TCF7L2* rs7903146 203, *BCL2* rs2279115 254

### Gene-environmental analyses

**Table 3** shows the interaction between NSAID and the polymorphisms. There was an interaction between use of NSAID and the *TP53* rs1042522 polymorphism (p_interaction_=0.04). *TP53* rs1042522 GG homozygotes had a lower relative risk of CRC for NSAID users to non-users compared to variant C-allele carriers.

**Table 3.** Interactions between polymorphisms and use of non-steroid anti-inflammatory drugs (NSAID)

|  | N<sub>cases</sub>/N<sub>sub-cohort</sub> | N<sub>cases</sub>/N<sub>sub-cohort</sub> | IRR crude (95%CI)<sup>1</sup> |  | IRR (95%CI)<sup>2</sup> |  |  |
| --- | --- | --- | --- | --- | --- | --- | --- |
|  | No | Yes | No | Yes | No | Yes | P-value<sup>3</sup> |
| SLC25A20 rs7623023 |  |  |  |  |  |  |  |
| AA | 241/469 | 120/216 | 1.00 | 1.09 (0.83-1.45) | 1.00 | 1.07 (0.80-1.42) |  |
| GA+GG | 397/717 | 159/320 | 1.08 (0.88-1.32) | 1.04 (0.81-1.34) | 1.09 (0.88-1.33) | 1.04 (0.81-1.35) | 0.59 |
| PRKAB1 rs4213 |  |  |  |  |  |  |  |
| TT+TG | 585/1077 | 246/489 | 1.00 | 0.97 (0.81-1.17) | 1.00 | 0.97 (0.80-1.17) |  |
| GG | 54/ 104 | 34/ 48 | 0.98 (0.69-1.39) | 1.32 (0.83-2.12) | 1.05 (0.74-1.50) | 1.39 (0.86-2.23) | 0.32 |
| LPCAT1 rs7737692 |  |  |  |  |  |  |  |
| AA+GA | 556/1032 | 241/473 | 1.00 | 0.99 (0.82-1.20) | 1.00 | 0.97 (0.80-1.18) |  |
| GG | 80/ 148 | 40/ 62 | 1.07 (0.79-1.44) | 1.34 (0.88-2.06) | 1.06 (0.78-1.44) | 1.41 (0.92-2.17) | 0.26 |
| PLA2G4A rs4402086 |  |  |  |  |  |  |  |
| AA+GA | 571/1091 | 253/491 | 1.00 | 1.04 (0.86-1.25) | 1.00 | 1.03 (0.85-1.25) |  |
| GG | 59/95 | 26/ 42 | 1.22 (0.86-1.73) | 1.22 (0.73-2.05) | 1.25 (0.88-1.77) | 1.20 (0.70-2.05) | 0.83 |
| ALOX5 rs3780894 |  |  |  |  |  |  |  |
| AA+GA | 616/1157 | 271/528 | 1.00 | 1.01 (0.84-1.21) | 1.00 | 1.00 (0.83-1.20) |  |
| GG | 16/29 | 8/9 | 1.07 (0.57-2.02) | 1.78 (0.66-4.74) | 1.06 (0.55-2.03) | 1.91 (0.71-5.11) | 0.33 |
| PTGER3 rs6685546 |  |  |  |  |  |  |  |
| TT+TC | 617/1151 | 273/520 | 1.00 | 1.01 (0.85-1.21) | 1.00 | 1.01 (0.84-1.21) |  |
| CC | 20/34 | 7/13 | 0.87 (0.48-1.55) | 1.05 (0.41-2.73) | 0.88 (0.49-1.60) | 1.03 (0.40-2.66) | 0.79 |
| TP53 rs1042522 |  |  |  |  |  |  |  |
| GG | 358/632 | 145/308 | 1.00 | 0.86 (0.68-1.09) | 1.00 | 0.85 (0.66-1.08) |  |
| GC+CC | 272/549 | 136/228 | 0.87 (0.72-1.07) | 1.11 (0.86-1.44) | 0.87 (0.71-1.07) | 1.11 (0.85-1.44) | 0.04 |
| CCAT2 rs6983267 |  |  |  |  |  |  |  |
| GG | 220/318 | 86/152 | 1.00 | 0.88 (0.64-1.21) | 1.00 | 0.87 (0.62-1.20) |  |
| TG+TT | 411/862 | 190/383 | 0.69 (0.55-0.85) | 0.74 (0.57-0.95) | 0.67 (0.54-0.83) | 0.71 (0.55-0.92) | 0.31 |
| TCF7L2 rs7903146 |  |  |  |  |  |  |  |
| CC+TC | 585/1104 | 251/498 | 1.00 | 1.00 (0.83-1.20) | 1.00 | 0.98 (0.81-1.19) |  |
| TT | 45/82 | 27/ 33 | 1.09 (0.74-1.60) | 1.63 (0.95-2.79) | 1.05 (0.71-1.55) | 1.64 (0.95-2.84) | 0.19 |
| BCL2 rs2279115 |  |  |  |  |  |  |  |
| AA | 181/339 | 87/159 | 1.00 | 1.13 (0.82-1.56) | 1.00 | 1.07 (0.77-1.48) |  |
| CA+CC | 427/833 | 183/364 | 0.95 (0.76-1.18) | 0.96 (0.74-1.24) | 0.91 (0.73-1.14) | 0.93 (0.71-1.21) | 0.82 |

IRR, incidence rate ratio; CI, confidence interval; BMI, body mass index; NSAID, non-steroidal anti-inflammatory drug; HRT, hormone replacement therapy.
<sup>1</sup>IRRs for CRC estimated by the Cox proportional hazards model with age as the underlying time axis, and stratified by gender, so that the underlying hazards are gender specific. 95% CI is based on Wald's tests.
<sup>2</sup>In addition, adjusted for smoking status, alcohol, HRT status (women only), BMI, use of NSAID, energy consumption, intake of red and processed meat, dietary fiber, intake of fruit and vegetable.
<sup>3</sup>P-value for interaction on a multiplicative scale
Number of missing values; *SLC25A20* rs7623023 254, *PRKAB1* rs4213 257, *LPCAT1* rs7737692 262, *PLA2G4A* rs4402086 266, *ALOX5* rs3780894 259, *PTGER3* rs6685546 259, *TP53* rs1042522 266, *CCAT2* rs6983267 272, *TCF7L2* rs7903146 268, *BCL2* rs2279115 320.

**Table 4** shows the interaction between dietary factors and the polymorphisms. Intake of red and processed meat interacted with *CCAT2* rs6983267 (p_interaction_=0.04). *CCAT2* rs6983267 T-allele carriers had a lower relative risk of CRC by meat intake compared to GG homozygotes. Furthermore, use of alcohol interacted with *TP53* rs1042522 (p_interaction_=0.04). The variant C-allele carriers increased their risk for CRC with increased alcohol intake wheres GG homozygotes did not. In the tertile analyses (**Supplemental Table 1**), *TP53* rs1042522 and *LPCAT1* rs7737392 variant allele carriers had a higher risk increase than GG homozygotes (p_interaction_=0.04 and 0.02, respectively). Furthermore, *SLC25A20* rs7623023 AA homozygotes had a higher risk increase than the variant G-carriers (p_interaction_=0.03) with increased meat intake. Variant allele carriers were at increased risk of CRC irrespectively of meat intake compared to the AA homozygotes. No other statistically significant interactions between diet or NSAID and the polymorphisms were found.

**Table 4.** Interactions between polymorphisms and dietary factors.

|  | IRR (95% CI)<sup>1</sup><br>Red and processed<br>meat (25 g/day) | P-value<sup>2</sup> | IRR (95% CI)<sup>1</sup><br>Fibre (10 g/day) | P-value<sup>2</sup> | IRR (95% CI)<sup>1</sup><br>Fruit and vegetables<br>(50 g/day) | P-value<sup>2</sup> | IRR (95% CI)<sup>1</sup><br>Alcohol (10<br>g/day) | P-value<sup>2</sup> |
| --- | --- | --- | --- | --- | --- | --- | --- | --- |
| *SLC25A20*<br>*rs7623023* |  |  |  |  |  |  |  |  |
| AA | 1.02(0.96-1.08) | 0.64 | 0.87(0.64-1.18) | 0.85 | 0.98(0.95-1.02) | 0.60 | 1.02(0.95-1.08) | 0.76 |
| GA+GG | 1.00(0.95-1.06) |  | 0.85(0.64-1.13) |  | 0.99(0.96-1.03) |  | 1.03(0.97-1.09) |  |
| *PRKAB1* *rs4213* |  |  |  |  |  |  |  |  |
| TT+TG | 1.00(0.95-1.05) | 0.45 | 0.85(0.65-1.12) | 0.11 | 0.99(0.96-1.02) | 0.30 | 1.02(0.97-1.07) | 0.60 |
| GG | 1.05(0.93-1.18) |  | 0.60(0.37-0.98) |  | 0.96(0.90-1.02) |  | 1.05(0.95-1.15) |  |
| *LPCAT1* *rs7737692* |  |  |  |  |  |  |  |  |
| AA+GA | 1.02(0.97-1.07) | 0.06 | 0.88(0.67-1.16) | 0.09 | 0.99(0.96-1.02) | 0.65 | 1.02(0.97-1.07) | 0.87 |
| GG | 0.92(0.84-1.02) |  | 0.65(0.43-0.98) |  | 0.98(0.93-1.03) |  | 1.03(0.92-1.15) |  |
| *PLA2G4A*<br>*rs4402086* |  |  |  |  |  |  |  |  |
| AA+GA | 1.01(0.96-1.06) | 0.66 | 0.88(0.67-1.16) | 0.92 | 0.99(0.96-1.02) | 0.92 | 1.03(0.98-1.08) | 0.74 |
| GG | 1.03(0.95-1.12) |  | 0.90(0.55-1.48) |  | 0.99(0.92-1.06) |  | 1.01(0.89-1.14) |  |
| *ALOX5* *rs3780894* |  |  |  |  |  |  |  |  |
| AA+GA | 1.01(0.96-1.06) | 0.65 | 0.86(0.65-1.13) | 0.85 | 0.99(0.96-1.02) | 0.47 | 1.02(0.98-1.07) | 0.88 |
| GG | 1.06(0.85-1.32) |  | 0.80(0.39-1.67) |  | 0.96(0.87-1.05) |  | 1.07(0.65-1.75) |  |
| *PTGER3* *rs6685546* |  |  |  |  |  |  |  |  |
| TT+TC | 1.01(0.96-1.06) | 0.23 | 0.85(0.65-1.11) | 0.85 | 0.99(0.96-1.02) | 0.90 | 1.02(0.98-1.07) | 0.64 |
| CC | 0.91(0.77-1.08) |  | 0.90(0.50-1.62) |  | 0.99(0.91-1.08) |  | 1.08(0.87-1.34) |  |
| *TP53* *rs1042522* |  |  |  |  |  |  |  |  |
| GG | 1.00(0.94-1.06) | 0.34 | 0.82(0.62-1.09) | 0.31 | 1.00(0.96-1.03) | 0.31 | 0.99(0.94-1.05) | 0.04 |
| GC+CC | 1.03(0.97-1.09) |  | 0.93(0.68-1.27) |  | 0.98(0.94-1.01) |  | 1.08(1.01-1.16) |  |
| *CCAT2* *rs6983267* |  |  |  |  |  |  |  |  |
| GG | 1.05(0.98-1.13) | 0.04 | 0.83(0.61-1.13) | 0.83 | 1.00(0.96-1.04) | 0.46 | 1.05(0.98-1.13) | 0.34 |
| TG+TT | 0.98(0.93-1.03) |  | 0.81(0.60-1.08) |  | 0.98(0.95-1.01) |  | 1.01(0.96-1.07) |  |
| *TCF7L2* *rs7903146* |  |  |  |  |  |  |  |  |

## Discussion

This large prospective investigated potential associations between polymorphisms in the fatty acid metabolic and NSAID pathways, and risk of CRC and, furthermore, the potential interaction between these polymorphisms and NSAID and diet (intake of red and processed meat, fiber, fruit and vegetables, and alcohol) in relation to CRC. The polymorphisms were selected from recent reviews based on their potential role in the fatty acid metabolic and NSAID pathways (**Table 5**) [22-24, 32]. We found that *CCAT2* rs6983267 GG genotype was associated with lowered risk of CRC per se and we found an interaction between the polymorphism and meat in relation to CRC. Furthermore, interactions between *TP53* rs1042522 and use of NSAID, alcohol intake, and, in the tertile analysis, intake of red and processed meat were found. Next, we found interactions between *LPCAT1* rs7737692 and *SLC25A20* rs7623023 and intake of red and processed meat in the tertile analysis in relation to CRC. No other consistent associations or interactions were found.

**Table 5.** uggested biological effects of the selected polymorphisms.

| Expected interaction | SNP ID | Nearby<br>gene | Allele | MAF | Bio effect | Ref |
| --- | --- | --- | --- | --- | --- | --- |
| Meat | rs7623023 | *SLC25A20* | G/A | 0.34 | Carnitine acylcarnitine translocase | [32] |
| Meat | rs4213 | *PRKAB1* | G/T | 0.31 | AMP-activated protein kinase β1 subunit | - |
| Meat | rs7737692 | *LPCAT1* | G/A | 0.36 | Lysophosphatidylcholine acetyltransferase | - |
| Meat | rs4402086 | *PLA2G4A* | G/A | 0.26 | Phospholipase A2 | - |
| Meat | rs3780894 | *ALOX5* | G/A | 0.16 | Arachidonate 5-lipoxygenase | - |
| Meat | rs6685546 | *PTGER3* | C/T | 0.14 | Prostaglandin E receptor 3 | - |
| Aspirin | rs1042522 | *TP53* | C/G | 0.46 | G allele increase p53 level | [36, 37] |
| Aspirin | rs6983267 | *CCAT2* | G/T | 0.39 | Aspirin suppresses the binding of TCF7L2 to the T allele | [22, 38] |
| Aspirin | rs7903146 | *TCF7L2* | T/C | 0.23 | Intron, transcription factor that plays a key role in the Wnt signaling pathway | [22] |
| Aspirin | rs2279115 | *BCL2* | G/F | 0.46 | Expression of BCL2 alternative splicing transcripts (BCL2-α, BCL2-β) in healthy donors | [36]<br>[39] |
| MAF, minor allele frequency; rs, reference SNP ID; SNP, single nucleotide polymorphism |  |  |  |  |  |  |

First, the association of *CCAT2* rs6983267 with CRC confirmed earlier results from several independent populations [25, 40] supporting the importance of the 8q24.21 gene locus for CRC carcinogenesis. The *CCAT2* rs6983267 polymorphism is located in a non-protein coding region near the *MYC* gene. The T-allele of *CCAT2* rs6983267 has been shown to impair binding of WNT/CTNNB1 pathway-related transcription factor 7 like-2 to DNA, thereby reducing *MYC* expression which in turn induces resistance to intestinal tumorigenesis [25]. The polymorphism has also previously been found to interact with aspirin. Nan et. al. found that variant T-allele carriers had 39-48% lower risk of CRC while using aspirin [25]. T-allele carriers of *CCAT2* rs6983267 constitute 27% of the sub-cohort members in the present study. As we did not find an interaction between *CCAT2* rs6983267 and NSAID use in the present study, the result may potentially suggest a specific effect of aspirin that may not be shared with non-aspirin NSAIDs in general. Unfortunately, the present study did not have the power to investigate aspirin use only.

Next, we found an interaction between *TP53* rs1042522 and NSAID. In our study, GG homozygotes lowered their risk of CRC by use of NSAID whereas variant C-allele carriers increased their risk of CRC by NSAID use (p=0.04). This is a replication of an earlier finding [41]. Tan et al., observed that GG homozygotes benefitted more from the use of NSAID than variant C-allele carriers. They found a substantial protective effect of NSAID use for homozygous carriage of the 72Arg allele compared to the 72Pro allele (odds ratio 0.44; 95% CI: 0.30–0.65) [41].

In the present study, 4 polymorphisms (*CCAT2* rs6983267, *TP53* rs1042522, *LPCAT1* rs7737692, and *SLC25A20* rs7623023) were found to interact with meat intake (**Table 4 and the S1 Table 1**). Two of the polymorphisms (*SLC25A20* rs7623023 and *LPCAT1* rs7737692) are involved in the metabolisms of fatty acids (**Table 5**), however, the functionality of the two common polymorphisms is unknown. The protein coded by *LPCAT1* is involved in the remodeling of phospholipids and has been associated with risk of sudden cardiac arrest [32], whereas the protein coded by *SLC25A20* is involved in the transport of fatty acids across the mitochondrial membrane. Our results may suggest that the fat from red and processed meat (that is metabolized to fatty acids) may contribute to the carcinogenic mechanism of red and processed meat in relation to CRC.

The two other polymorphisms (*CCAT2* rs6983267 and *TP53* rs1042522) have been found to interact with aspirin/NSAID in relation to CRC in the present or other studies [25, 41]. *TP53* rs1042522 is a missense polymorphism in the *TP53* gene where Arginine is changed to Proline, which results in increased apoptosis potential due to increased p53 levels [36, 37]. Several epidemiological studies, including randomized controlled clinical trials, have demonstrated that NSAID use decreases the incidence of adenomatous polyps and CRC [5]. The mechanism is thought to be caused by cell-cycle regulation and/or induction of apoptosis via mechanisms dependent and independent of cyclooxygenase [5, 42]. The use of NSAID may enhance the apoptosis potential already present in the GG genotype of *TP53* rs1042522 resulting in decreased risk of CRC compared to variant C-carriers. A diet high in meat was associated with increased risk of CRC among variant C-allele carriers compared to those with a diet low in meat intake. We have previously shown that intake of meat interacts with polymorphisms in inflammatory genes in relation to CRC risk [17, 18, 35] suggesting that a diet high in meat may cause an inflammatory milieu that increases the carcinogenic potential in persons with an impaired *TP53* gene. This hypothesis could also apply for the *CCAT2* rs6983267 polymorphism since persons homozygous for the G-allele have a higher expression of *MYC* [38] and thereby an increased carcinogenic potential which could be further triggered by a diet high in meat. The finding that alcohol intake interacted with *TP53* rs1042522 resulting in increased risk of CRC for variant C-carriers may be caused by a similar mechanism as meat since alcohol is known to be associated with a systemic inflammatory state [43] and thus the protective effect of the G-allele is abolished.

Advantages and limitations with the study design have been described in previous studies [15-21]. The main advantage of this study is the prospective study design with collection of dietary and lifestyle factors before diagnosis that eliminates the risk of recall bias. Another main advantage is the diverse and high intake of meat in the present cohort enabling identification of gene-meat interactions. The prospective “Diet, Cancer and Health” cohort has proven to be suitable to detect meat-gene interactions [17, 18, 35]. Changes in dietary and lifestyle habits during follow-up is possible but, if present, will result in lower power to detect real differences between cases and the comparison group. The “Diet, Cancer and Health” cohort is homogenous reducing population specific genetics and dietary patterns seen in larger multicentre studies. The disadvantage of the prospective study is the limited power to study gene-environment interactions. None of the results withstood Bonferroni correction. Thus, all new findings should be sought replicated in independent prospective cohorts with well-characterized lifestyle information.

## Conclusions

In conclusion, in this study an association of *CCAT2* rs6983267 with CRC and an interaction between *TP53* rs1042522 and NSAID in relation to CRC were replicated. Our exploratory analyses found interactions between polymorphisms in the fatty acid metabolic pathway (*LPCAT1* s7737692 and *SLC25A20* rs7623023) and polymorphisms that have been found to interact with NSAID/aspirin (*CCAT2* rs6983267 and *TP53* rs1042522) on one hand and intake of red and processed meat on the other in relation to risk of CRC. Our results suggest that meat intake and NSAID use affect the same carcinogenic mechanisms. All new findings from this study should be sought replicated in independent prospective cohorts with well-characterized lifestyle information. Future studies on the cancer-protective effects of aspirin/NSAID should include gene and meat assessments.

## Materials and Methods

### Subjects

As previously described [26] the “Diet, Cancer and Health” Study is an ongoing Danish cohort study designed to investigate the relation between diet, lifestyle and cancer risk. The cohort consists of 57,053 persons, recruited between December 1993 and May 1997. All the subjects were born in Denmark, and the individuals were 50 to 64 years of age and had no previous cancers at study entry. Blood samples, anthropometric measures and questionnaire data on diet and lifestyle were collected at study entry.

### Follow-up and endpoints

As previously described [20] the present study used a nested case-cohort design. Follow-up was based on population-based cancer registries. Between 1994 and 31st December 2009, 1038 CRC cases were diagnosed. A sub-cohort of 1857 persons was randomly selected within the full cohort at the time of entry into the cohort in agreement with the case-cohort study design [27] and, thus, without respect to time and disease status. Due to the study design, with a priori sampling of the sub-cohort, 28 persons were both cases and sub-cohort, and these persons were kept in the analyses. All 1038 CRC cases and 1857 sub-cohort members were included in the analysis. Flowchart of the participants is shown in the **S1 Fig 1**.

### Dietary and lifestyle questionnaire

Information on diet, lifestyle, weight, height, medical treatment, environmental exposures, and other socio-economic factors were collected at enrolment using questionnaires and interviews and has been described in details elsewhere [20, 28]. In short, the food-frequency questionnaire, assessed diet consumption in 12 categories of predefined responses, ranking from ‘never’ to ‘eight times or more per day’. The daily intake was then calculated by using FoodCalc [29]. Red meat was calculated by combining intake of fresh and minced beef, veal, pork, lamb, and offal, whereas processed meat combined intake of bacon, smoked or cooked ham, other cold cuts, salami, frankfurter, Cumberland sausage, and liver pâté. The total dietary fiber was estimated by the method of the Association of Official Analytical Chemists [30], which includes lignin and resistant starch. Fiber intake is calculated by multiplying the frequency of consumption of relevant foods (i.e. fruit, vegetables, grains, and leguminous fruit) by their fiber content as determined from national databases of food content. For fruit, only intake of fresh fruit was examined, whereas intake of vegetables also included estimated contributions from food recipes. Intake of alcohol was inferred from the food-frequency questionnaire and lifestyle questionnaire as described in details in [31]. Abstainers were defined as those who reported no intake of alcohol on the food-frequency questionnaire and no drinking occasions on the lifestyle questionnaire. Smoking status was classified as never, past or current. Persons smoking at least 1 cigarette daily during the last year were classified as smokers. NSAID use (“Aspirin”, “Ibuprofen”, or “Other pain relievers) was assessed as ≥ 2 pills per month during one year at baseline. Use of hormone replacement therapy among women was assessed as current, former or never user.

### Genotyping and selection of polymorphisms

The polymorphisms were chosen based on Andersen et. al. [22] and Lemaitre et. al. [32]. Promising polymorphisms with known functionality or that were associated with biological effects suggesting functionality or linkage with functional polymorphism and with a reasonable minor allele frequency to study gene-environment interactions were selected. Buffy coat preparations were stored at minus 150°C until use. DNA was extracted as described [33]. The DNA was genotyped by LGC KBioscience (LGC KBioscience, Hoddesdon, United Kingdom) by PCR-based KASP™ genotyping assay (lgcgenomics.com/). To confirm reproducibility, genotyping was repeated for 10% of the samples yielding 100% identity.

## Statistics

Incidence rate ratios (IRR) and 95% Confidence Interval (CI) were based on a Cox proportional hazard model fitted to the age at the event of CRC according to the principles for analysis of case-cohort [27] using the approach of Prentice and Langholz [34]. The main explanatory variables were the polymorphisms. All models were adjusted for baseline values of risk factors for CRC as published previously [17-21, 35]; body mass index (BMI) (kg/m 2, continuous), use of hormone replacement therapy, (never/past/current, among women), intake of dietary fibre (g/day, continuous), and processed red meat (g/day, continuous), energy intake (kJ/day), NSAID use (yes/no) and smoking status (never/past/current). Cereals, fiber, fruit, and vegetables were also entered linearly as continuous covariates. All analyses were stratified by gender so that the basic (underlying) hazards were gender specific.

In the interaction analyses of the dietary factors with polymorphisms, we present two analyses: in one analysis the dietary factors were used as numeric variables and in the other, they were entered in the models as a three-level categorical variable defined via tertile cutpoints derived from the empirical distribution of the whole population. Information on numbers of missing observations on lifestyle data and genetics are included the individual tables. In addition, for the interaction analyses, all abstainers of alcohol were excluded from the analyses. Deviation from Hardy-Weinberg equilibrium in the comparison group was assessed using a Chi-square test. All analyses were performed using the survival package (Terry M. Therneau, version 2.42.4) of the statistical computational environment R, version 3.5.1. A p<0.05 was considered to indicate a statistically significant test result.

## Ethics

All participants gave verbal and written informed consent. The Diet, Cancer and Health study was approved by the National Committee on Health Research Ethics (journal nr. (KF) 01-345/93) and the Danish Data Protection Agency.

## Acknowledgements

We thank Katja Boll for technical assistance with data managing. The project was supported by “Familien Erichsens Mindefond”, “Købmand Sven Hansen og hustru Ina Hansens Fond” and “Knud og Edith Eriksens Mindefond”. The funders had no role in study design, data collection, and analysis, decision to publish, or preparation of the manuscript.

## Additional Files

**S1 Fig 1. Flowchart of study participants.**

**S1 Table 1. Tertile analyses of polymorphisms and dietary factors.**

Author contributions to the manuscript
UH performed the statistical analyses, VA wrote the first draft of the manuscript. VA, TIK, and UV conceived the study, TIK and VA interpreted the data, critical revised the manuscript for important intellectual content and VA obtained funding. AT designed the cohort study and collected the biological material. All authors commented on the work and accepted the final manuscript.

## Abbreviations used in this manuscript

CI: confidence intervals
CRC: colorectal cancer
GxE: Gene-environment
IRR: Incidence rate ratios
NSAID: non-steroidal anti-inflammatory drug

